# Modulating Cellular Deformability via 3D Dextran Hydrogel Cultivation to Regulate the Microcirculation of Mesenchymal Stem Cells in Murine Spleen and Liver

**DOI:** 10.1101/2024.12.15.628544

**Authors:** Xiaolu Zhu, Zheng Wang, Yuanping Shi, Shuang Yao, Fengliang He, Fang Teng

## Abstract

For mesenchymal stem cell (MSC) therapy to be effective, the vascular system may be used to deliver and steer the cells to the target tissue. However, the expanded MSCs in petri dishes are usually have a limited deformability and commonly excluded by the capillary networks when homing to the downstream organs via microcirculation. Here, we propose to utilize specially designed 3D dextran hydrogels and tuning the microscopic heterogeneity of hydrogel composition to make the administrated cells mechanically comply with the structure and mechanics of the capillary. The cell deformability in petri dishes, microcosmically homogeneous (HOM) and heterogeneous (HET) dextran hydrogels were investigated in vitro by measuring cell moduli through atomic force microscope (AFM), analyzing the expression of cytoskeletal protein via flow cytometry and fluorescent imaging. The in vitro experimental results demonstrate a progressive increase in cell deformability from 2D dishes, to HOM-hydrogel derived cells, and then to HET-hydrogel derived cells. The in vivo mouse experiment indicates the cells could deform accordingly and pass through easily with reduced resistance inside the mouse organs. It is suggested that the main destination of hMSC microcirculation could be selected between the spleen and liver of mice, by tuning cell mechanics that depends on the stimulus from microcosmically HOM or HET hydrogel, which lays a potential foundation for the mechanically modified MSC therapy for lesions in the organs.

## 1. Introduction

*In vitro* cell culture technologies provide a versatile platform to manufacture cell products with desired characteristics(1), such as controllable migration dynamics(2, 3), maturated differentiation markers(4), enhance paracrine signaling (5, 6). As the production of various modified cell types proliferates, the challenges of their rational application in medicine have become increasingly important and pressing (7-9). Notably, the mechanical properties of cells have been shown to have intricate connection with their physiological and pathological behaviors (10-14). This relationship has critical biomedical applications, including mechanical modeling of tissue morphogenesis (10, 15, 16), pathological diagnostics of tissue cirrhosis and fibrosis (17), and mechanisms of tumor tissues(18) and tumor cell invasion (19), cellular mechanics in the context of disease progression and therapeutic interventions(20, 21). In particular, the mechanical properties of stem cells, including their elasticity and viscoelasticity, play a crucial role in regulating their behavior, including proliferation, differentiation, and response to external stimuli (22). For instance, studies have shown that the mechanical environment, such as the stiffness of the extracellular matrix, can significantly influence stem cell fate decisions(23). Additionally, the cytoskeletal architecture and mechanical properties of stem cells are essential for their interaction with biomaterials, which are often used in regenerative medicine applications(24). However, the impacts of mechanical properties of stem cells on the delivery efficiency during therapeutic processes remain considerably less understood (11, 25).

Microcirculation plays a pivotal role in the delivery of therapeutic cells to target sites (26). Unfortunately, the efficiency of this delivery is often hindered by the physical barriers presented by pulmonary capillary networks (27). Cells can adapt to extracellular confinement via biomechanical adjustments, which is key for regulating cellular elasticity to travel through capillary networks. For example, with specifically tailored culture medium with serval supplements and growth factors, the cells cultured in ultralow attachment culture flasks formed spheroids of human Mesenchymal Stem Cells (hMSC) exhibited lower Young’s modulus and enhanced rheological behavior, compared to those of flat cells grown on 2D tissue culture polystyrene (TCPS) (26). The mesenspheres could also be formed by allocating MSCs in microwells on the hydrogel surface (28), or in hyaluronic acid/alginate (HA@Alg) core-shell microcapsules, aiming for stem cell-based therapies(29), yet these studies have not further investigated the mechanical modulus variation of MSCs derived from the formed multicellular spheroids.

Recent studies have demonstrated the feasibility of reducing cellular modulus utilizing 3D cell culture technique, which requires no specifically tailored culture medium or specially constraint structures, and increases the likelihood of cells passing through the blood capillaries with micron-level diameters (30), but the further demonstration by in vivo animal experiments still lacks for facilitating the delivery of therapeutic stem cells to organs such as the liver and spleen via standard intravenous injection. Moreover, the corresponding method to control the microcirculation destination of administrated cells is scarce, which is crucial in clinical treatments.

In this research, we investigate and validate the improved microcirculation performance of hMSCs in murine spleen and liver by modifying the degrees of cellular deformability via the stimuli from the engineered three-dimensional (3D) hydrogels with tunable component heterogeneity. Cellular deformability is modulated by culturing cells within the 3D dextran hydrogels containing crosslinking molecules distributed in optimized degrees of heterogeneity, with cells exhibiting high or moderate deformability referred to as HD-cells or MD-cells, respectively. Low deformability cells (LD-cells), serving as a control group, are cultured using tissue culture polystyrene (TCPS) dishes. Effectiveness of the fabricated hydrogels with tunable microcosmic heterogeneity on regulating cell deformability was validated by measuring cell moduli through atomic force microscope (AFM), analyzing the expression of cytoskeletal protein via flow cytometry and fluorescent imaging, and quantifying the human Alu sequences inside hMSCs for verifying the existence of hMSCs in the organs of mice through real-time quantitative polymerase chain reaction (RT-qPCR). Notably, animal experiments reveal that cells with enhanced deformability can traverse the lungs and effectively infiltrate the liver and spleen. Moreover, MD-cells showed a significantly higher retention in the spleen after passing through the lungs, while HD-cells predominantly accumulated in the liver. Collectively, these findings suggest that the microcirculation destination of hMSCs can be engineered through the manipulation of cell deformability in vivo, with promising implications for medical therapies.

## 2. Results

### 2.1 Characterization of 3D heterogeneous and homogeneous hydrogels

Heterogeneous hydrogels were fabricated by cross-linking the mixed polymer of dextran with different amount of available maleimide groups, based on thiol-maleimide addition reaction (Fig. 2). Cross-linking heterogeneity was regulated by tuning the amount ratio of two sets of available maleimides. The fabrication process of the 3D dextran heterogeneous hydrogels is depicted in Fig. 2A. Briefly, two equal-volume aliquots of maleimide-dextran, each functionalized with varying quantity of available cross-linking sites, were thoroughly mixed and fully cross-linked. The available quantities of maleimide group for the two aliquots are denoted as α and β. In this design, the α is not equal to β because the reactive maleimide groups on dextran macromolecules in the two aliquots were blocked using different quantities monothioglycerol via thiol-Michael addition. The concentrations of used monothioglycerol of the two aliquots were 0.67 mM and 0.29 mM respectively for heterogeneous (HET) hydrogel, and both were 0.48 mM for the homogeneous (HOM) hydrogels in which the α = β as indicated in Fig. 2B. Concentration values of the components for the HOM and HET hydrogels can be found in Table 1 in the section of materials and method. In the semi-synthesized hydrogel, this approach enhances the likelihood that two spatially adjacent dextran macromolecules form covalent bonds with markedly different quantities of the cross-linker, as schematically presented by the typical transections within the 3D hydrogel indicated in Fig. 2, a phenomenon referred to as the cross-linker clustering (CLC) effect by us.

**Table 1.**
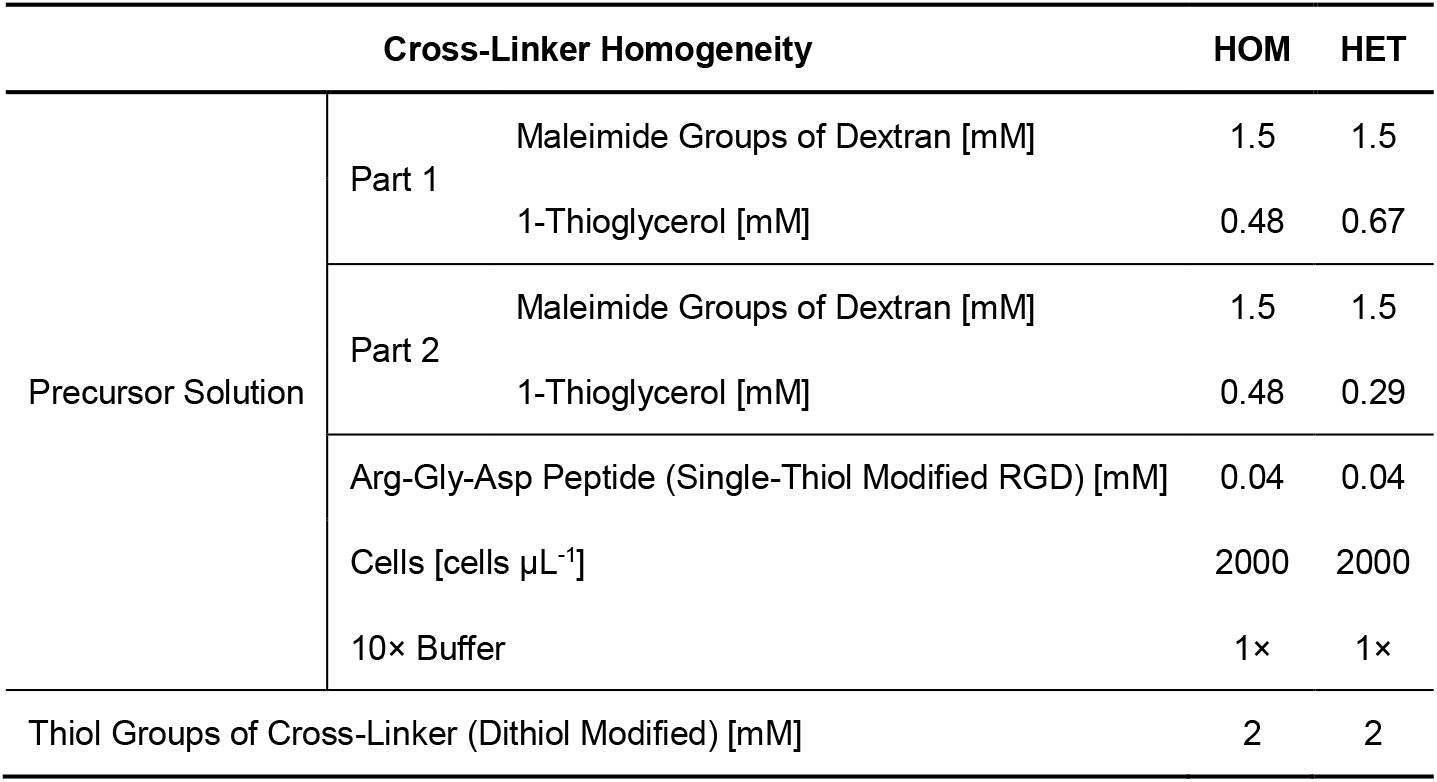
Final concentration of the components in formed hydrogel.

| Cross-Linker Homogeneity |  | HOM | HET |
| --- | --- | --- | --- |
| Precursor Solution | Part 1 Maleimide Groups of Dextran [mM] | 1.5 | 1.5 |
|  | 1-Thioglycerol [mM] | 0.48 | 0.67 |
|  | Part 2 Maleimide Groups of Dextran [mM] | 1.5 | 1.5 |
|  | 1-Thioglycerol [mM] | 0.48 | 0.29 |
|  | Arg-Gly-Asp Peptide (Single-Thiol Modified RGD) [mM] | 0.04 | 0.04 |
| | Cells [cells $\mu\text{L}^{-1}$] | 2000 | 2000 |
|  | 10× Buffer | 1× | 1× |
| Thiol Groups of Cross-Linker (Dithiol Modified) [mM] |  | 2 | 2 |

**Figure 1.**
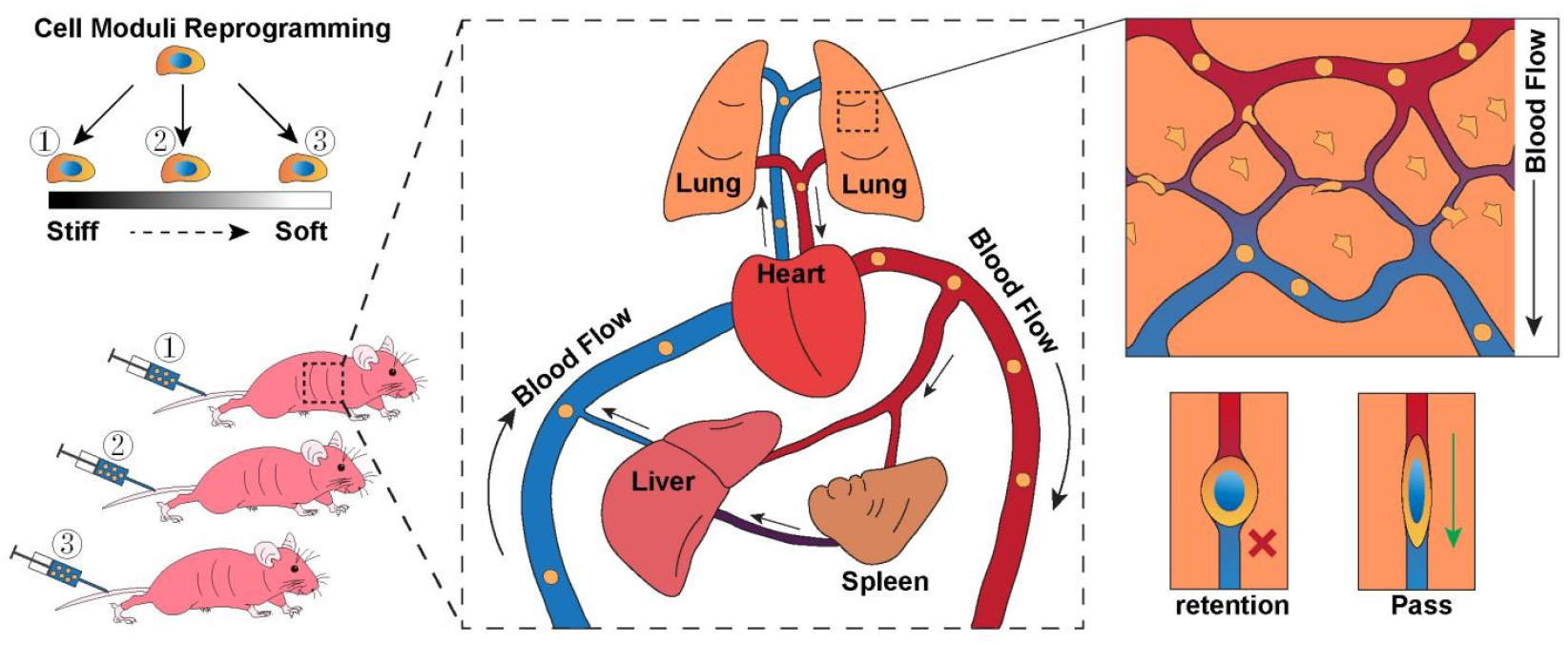
Schematically Illustration of hMSCs microcirculation *in vivo* after modification via *in vitro* culture.

**Figure 2.**
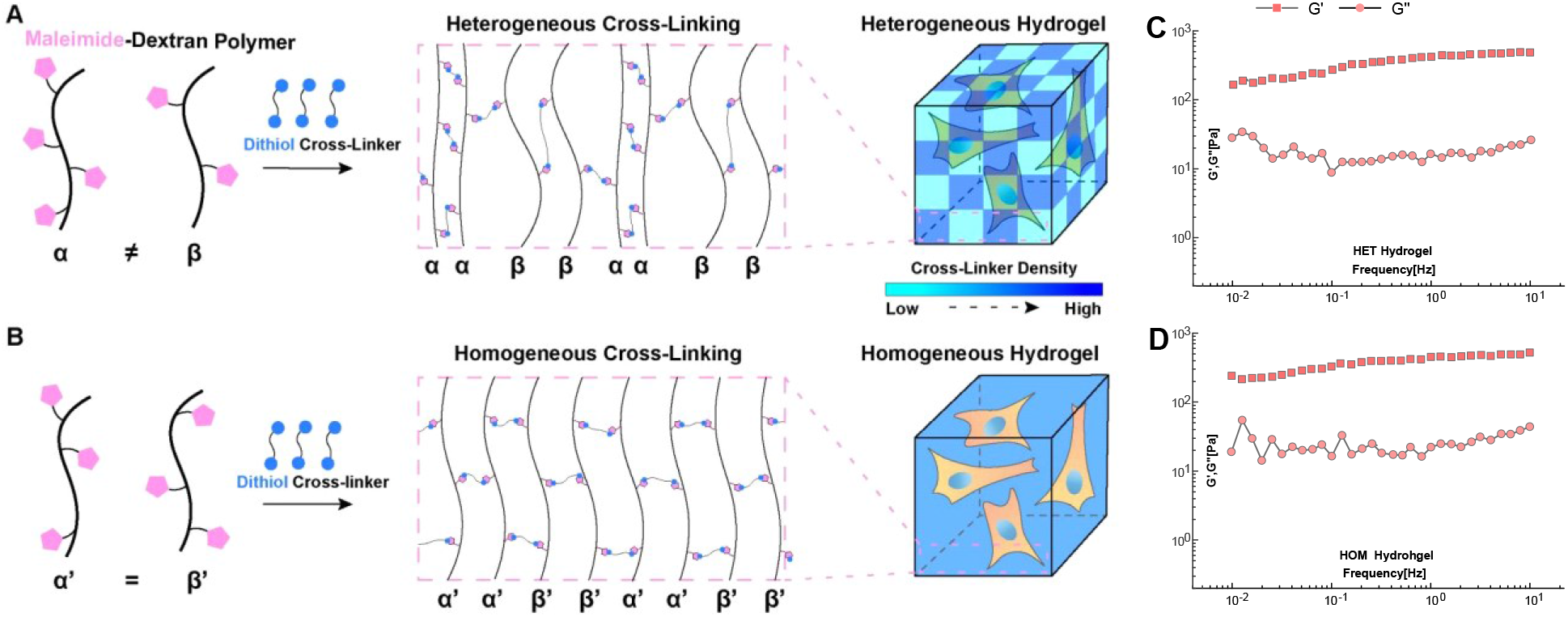
Schematically illustration of fabricating (A) heterogeneous, and (B) homogeneous 3D hydrogels and the characterization results of (C-D) macroscopic rheological properties of the hydrogels.

Rheological properties of microcosmically heterogeneous (HET) and homogeneous (HOM) hydrogels used in present study are basically comparable, which are averagely 341.8 and 373.7 Pa for storage moduli, and 17.6 and 25.8 Pa for the loss moduli, respectively, since the micro-sites with less crosslinking in HET hydrogel may let the elasticity and viscosity of the macroscopical hydrogel reduced than the HOM hydrogel. The elastic modulus (G′) of both HET and HOM hydrogels was over 10 times higher than the viscous modulus (G″) (without crossing points) for most of the frequency range of 0.01™10 Hz. Such fabrication strategy gives a heterogeneous distribution of local stiffness of hydrogels in microscales, and it possesses the controllable and appropriate viscoelasticity to provide the mechanical support for 3D culture of hMSCs.

### 2.2 Quantitatively Characterization of Deformability of MSCs Modified via 3D Hydrogel Culture

Young’s modulus of cells from two hydrogel groups and TCPS group were probed via AFM. Cells were extracted from hydrogels or TCPS surface, and placed on the glass plate for AFM measurement (Fig. 3A). The results showed the highest moduli value of cells from TCPS group, which indicated a most rigid cell body from TCPS group compared with other two hydrogel groups (Fig. 3B). Further, results of morphological analysis indicate the largest circularity and smallest averaged aspect ratio (AR) of cell nuclei in TCPS group compared with cells from hydrogel groups (Fig. 3C-D). It’s another complementary proof of low deformability of TCPS cells. Low deformability could significantly influence microcirculation of cells because nuclear deformation is necessary for cell to pass through the capillary with diameter of around 2-3 µm. In addition, AFM results showed that cells from HET hydrogel group had lower moduli than cells from HOM hydrogel group. It supported that stiffness-heterogeneity of 3D hydrogel decreased cell moduli and enhanced its deformability. Moreover, the statistic of the FCM data of F-actin are shown in Fig. 3E-F, and the result of averaged immunofluorescence analysis (Fig. 3G) indicates lower F-actin expression of HD cells compared with LD cells. It further supports the lower rigidity (also known as higher deformability) of the cells inside HET hydrogels, since intracellular F-actin density usually positively corresponds to cell rigidity. It could be reasonable to infer that lower F-actin density resulted in stronger fluidity of cells, which deformed more flexibly and probably passed through capillaries more easily during microcirculation. Additionally, the multicellular mesenspheres consisting of crowed MSCs usually can be formed inside the 3D hydrogels, and the spatial distributions of fluorescent intensity of F-actin are shown in Fig. 4. The fluctuation of fluorescent intensity is more significant in the HET-gel group which is possible due to the crosslinker clustering stimulates the reorganization of multicellular cytoskeletons inside mesenspheres. All these results demonstrate a successive ascent of potential deformability from TCPS cells (high moduli), to HOM-hydrogel derived cells (moderate moduli), and to HET-hydrogel derived cells (low moduli), which can be further applied in the investigation of correspondence between deformability of administrated cells and the microcirculation results.

**Figure 3.**
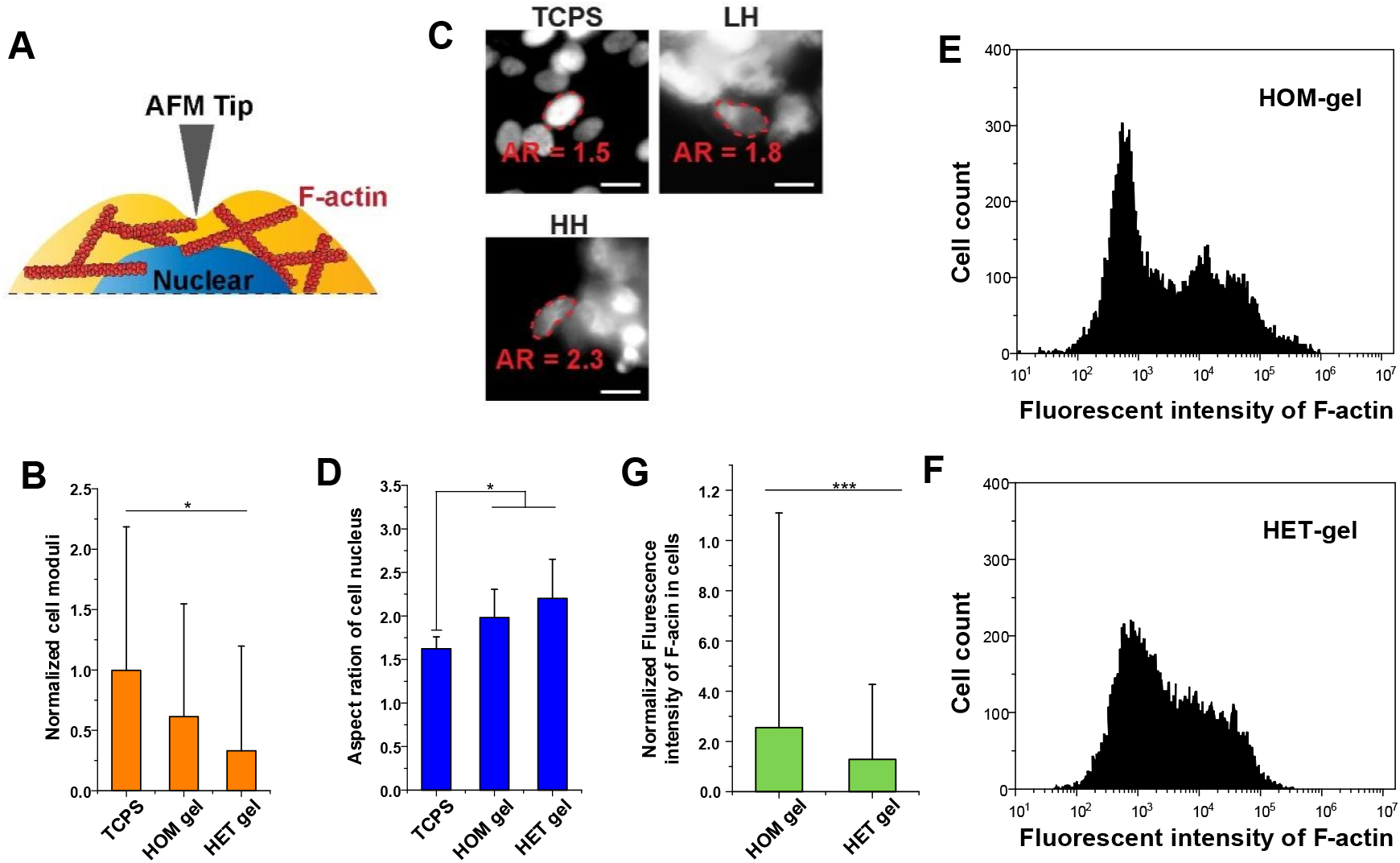
Characterization of cell deformability. (A) Illustration of probing cell moduli via atomic force microscopy (AFM); (B) Cell moduli measured via AFM was compared among groups of cells from tissue culture polystyrene (TCPS), homogeneous (HOM) hydrogels, and heterogenous (HET) hydrogels (n = 100 cells/group). (C) Images of representative cell nuclear morphology from each tested group; scale bar indicated 20 µm. (D) Statistical results of measured aspect ratio (AR) of cell nuclear (n = 60 cells/group). F-actin of cells from (E) HOM-gel group (n = 10393 cells), and (F) HET-gel group (n = 9515 cells) was quantified via flow cytometry. (G) Flow cytometry results of cellular F-actin from LH group (n = 10393 cells), and HH group (n = 9515 cells) were statistically analyzed. Quantitative data are represented as the mean ± SD. Statistical analysis was performed using the one-way ANOVA test and is represented: \**P* < 0.05, \*\*\**P* < 0.001.

**Figure 4.**
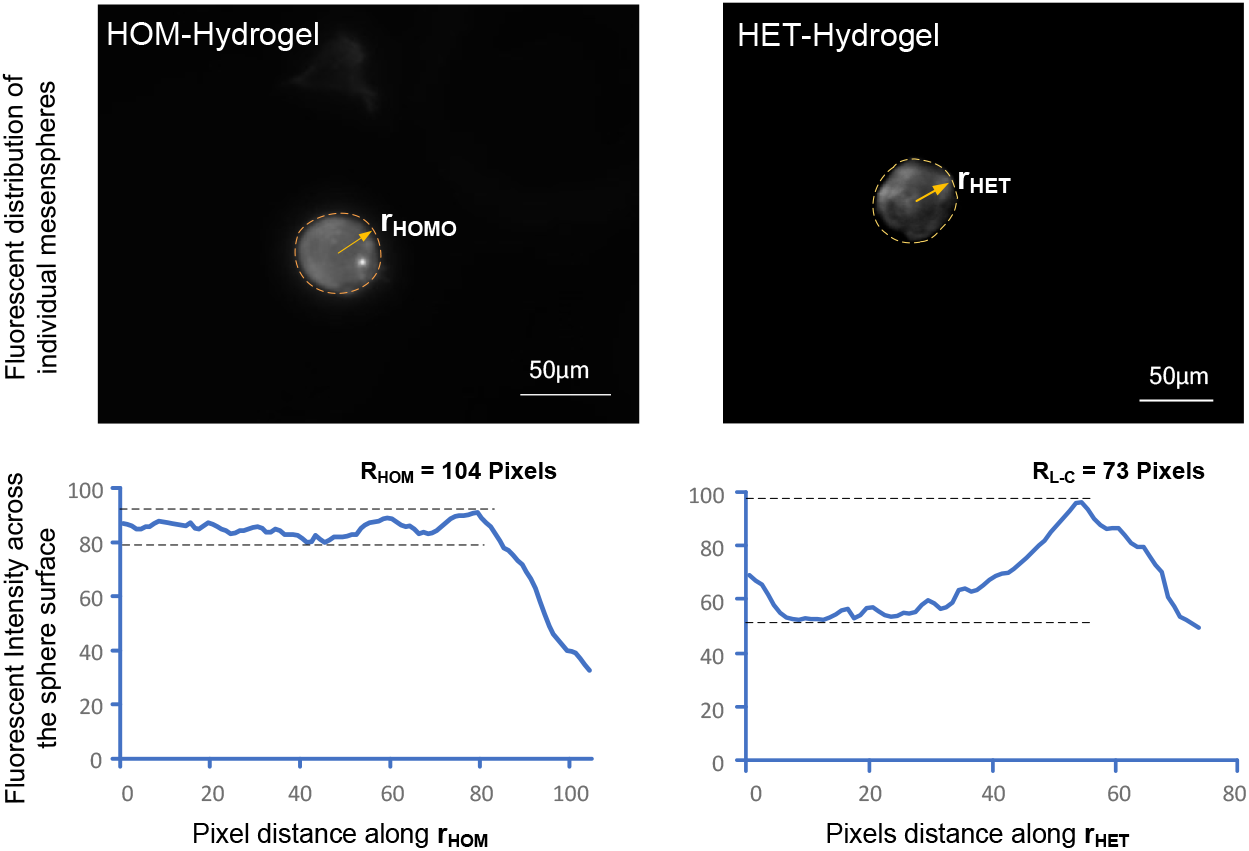
Characterization of fluorescent distribution of F-actin in typical multicellular mesenspheres.

### 2.3 The hMSCs amount retained in organs of lung, spleen and liver via improved microcirculation of hMSCs in mice

Next, the functions of the biomechanically modified hMSCs were evaluated *in vivo* to disclose their ability to bypass sequestration in the microcirculation. Resuspended hMSCs in normal saline were intravenously injected via the tail vein of mice, as illustrated in Fig. 5A. The delivered hMSCs were supposed to circulate in the blood capillary networks, forced by heart pumping (Fig. 1). The vital organs, including the lungs, spleens and livers were taken to check the amount of human cells able to locate into them from the circulation. Primers targeting the human ALU sequence were used to specifically identify and quantify human DNA via PCR analysis, which is a highly sensitive and specific quantification method for human cells (31). Compared with moderately and highly deformable groups, significantly high amount of hMSCs DNA were detected in lungs in low deformable group (petri dish group) as shown in Fig. 5B. It indicates that hMSCs were retained in lungs in TCPS group, while cells in the other two groups could pass through the lungs and get into livers and spleens more effectively. It supported that rigid cells were easily captured and retained in pulmonary capillary networks. Meanwhile, the moderately deformable group indicated markedly high DNA amount in spleen, as compared with the other two groups (Fig. 5C). Firstly, it implied that cells with stronger deformability could pass pulmonary capillary networks more likely, when cells with low deformability were apt to retain in lungs. Secondly, cells from moderately deformable group might have more chances to stay in spleen capillary networks. Lastly, cells with high-level deformability showed highest DNA amount in livers compared with the other two groups (Fig. 5D). Similarly, it supports that high deformable cells bypass the limitation of pulmonary capillary networks. Consequently, these results suggested that highly deformable cells were more capable to stay in liver capillary networks, instead of staying in spleens. Therefore, we supported that the retention ratio of administrated cells in different organs (lung, spleen, liver) corresponded with the levels of cellular deformability. This could be applied to improve the retention ratio of hMSCs in target organs during therapeutic microcirculation.

**Figure 5.**
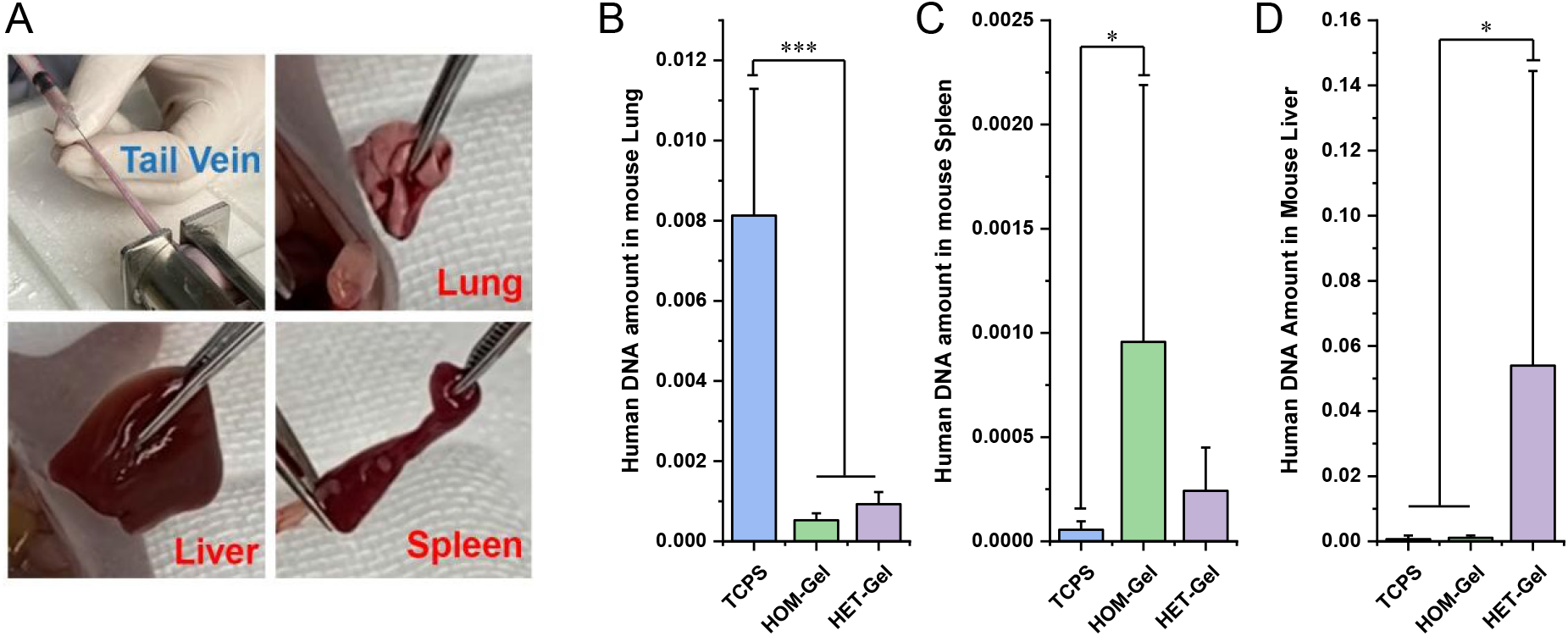
The hMSCs amount in lungs, spleens and livers of mice after injecting three kinds of deformable cells into mice. (A) The hMSCs in saline were injected into the nude mice via tail vein. After 20 min, the lungs, spleens, and livers were taken to detect the relative quantity of human DNA. The relative quantity of human DNA in the mouse organs was analyzed by the real-time quantitative polymerase chain reaction (RT-qPCR): (B) lungs, (C) spleen; (D) liver (n = 12 tests/group). Quantitative data are represented as the mean ± SD. Statistical analysis was performed using the one-way ANOVA test and is represented: *P < 0.05, ***P < 0.001.

The transmission electron microscopy (TEM) images of the lumens of capillary vessels within mouse organs (lung, spleen and liver), following administration of hMSCs, are presented in Fig. 6. The presence of exogenous cell structures was observed within the cross-sections of capillary vessels whose short diameters ranging from 3 to 5 microns in spleen and liver, and more than 5 microns in the lung. This observation confirms the retention of hMSCs in mouse organs. The diameter of MSCs decreased to less than 3 microns in the spleen and even less than 2 microns in the liver, likely due to the rapid flushing and squeezing by blood fluid.

**Figure 6.**
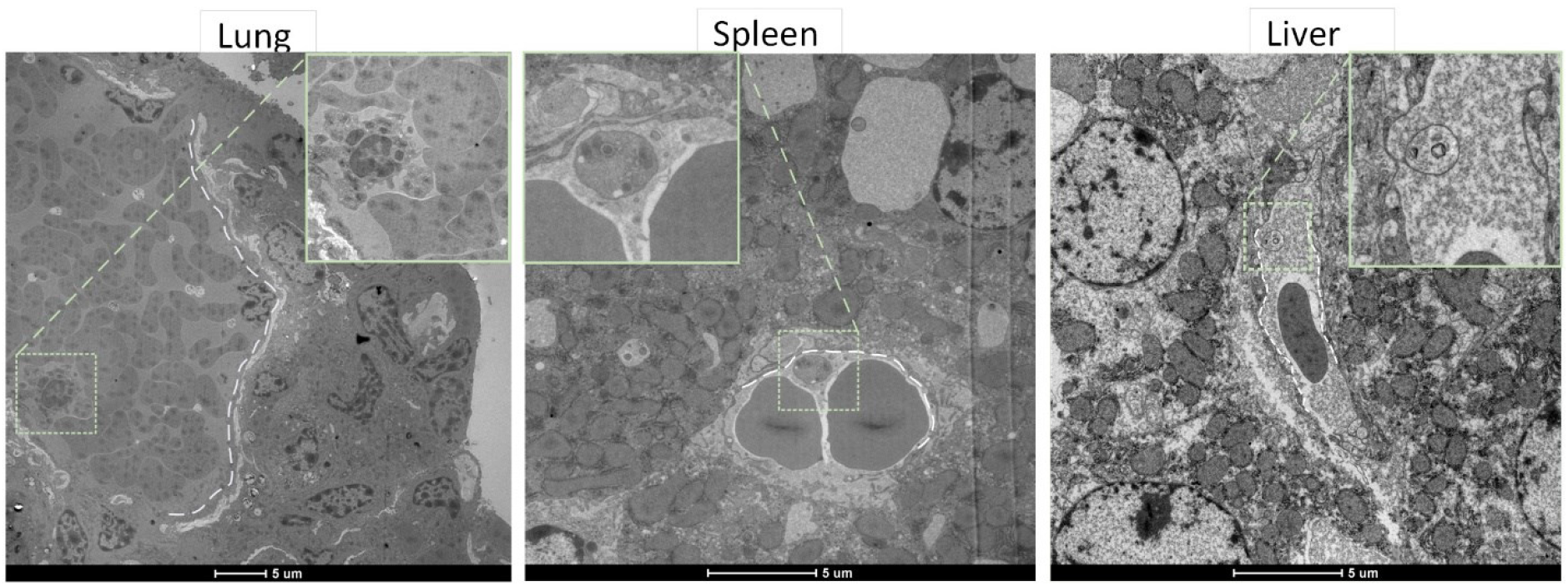
The TEM images of the lumens of capillary vessels inside mouse organs (lung, spleen and liver) after the organ samples were taken, fixed and sliced. The ultra-structures of the capillary lumens and surrounding microtissues in mouse lung, spleen and liver were obtained. The wireframe boxes indicate the location of the modified hMSCs administrated into capillaries.

## 3. Discussions

3D substrate-based cell culture opens a new world for reprogramming cellular properties *in vitro* in a more mimetic manner, resulting in a booming research of manufacturing cells with therapeutic effects. To examine their effectiveness, operable methods of delivering therapeutic cells into an *in vivo* target site precisely are urgently needed. This study regulated the deformability of hMSCs via 3D stiffness-heterogeneous hydrogel culture, and examined their performance in microcirculation test. Attentions were mainly focused on the microcirculation of hMSCs in heart-lung-spleen-liver circuit as shown in Fig. 7.

**Figure 7.**
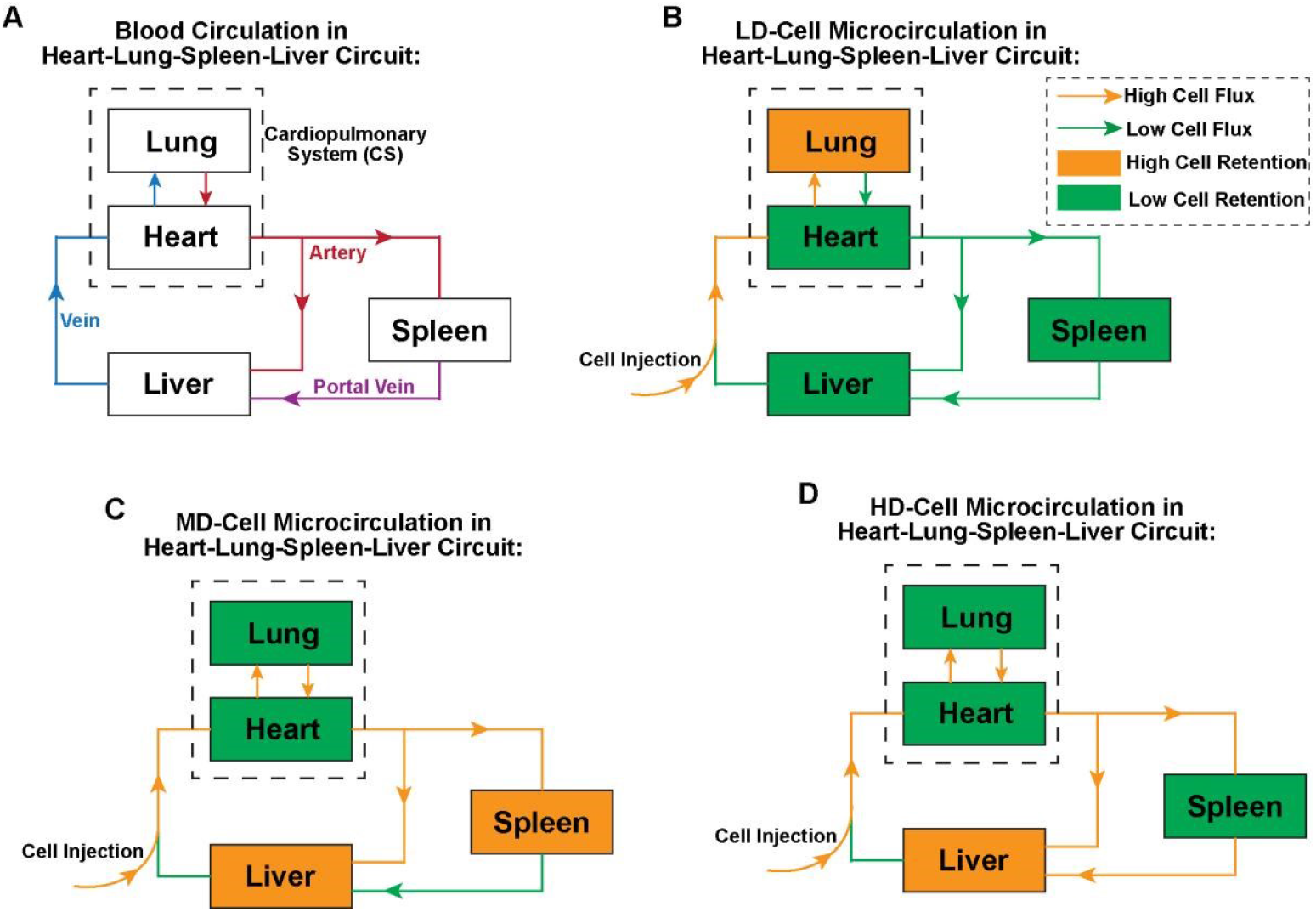
Illustration of hMSCs microcirculation in heart-lung-spleen-liver circuit. (A) Heart, lung, spleen, liver form a circulation for blood flow. Microcirculation tracks of cells with (B) low deformability (LD), (C) moderate deformability (MD), and (D) high deformability (HD) were illustrated.

Each organ is filled with abundant capillary networks, scaled with 2–5 µm inner diameter, for blood delivery. Previous, quite amount of reports indicated that TCPS derived cells can easily be trapped in pulmonary tissues, which is consistent with result of the TCPS group in the present study. This is a two-side coin. On one hand, it can be directly applied in cell therapies for pulmonary disease treatment, because pulmonary capillary serves as the natural trapper and collector of the administrated therapeutic cells. On the other hand, it could limit the extended application of cell microcirculation into other organs like heart, liver, spleen, kidney, brain and so on. One possible solution to the latter problem is to make the administrated cells mechanically comply with the structure and mechanics of the capillaries, so that cells could deform accordingly and pass through easily with reduced resistance. The hMSCs with lower modulus showed more DNAs retained in spleen or liver, instead of lung. That elucidates that softer or more deformable cells are more likely to bypass lung and get into the other organs. Benefit from this, microcirculation-based cell therapies would possibly be further applied in treating diseases in other organs. Meanwhile, it is followed by another question that whether the effect of different deformability of cells on microcirculation in other organs has the same law like that in lung. Our results showed that hMSCs with strongest deformability were most likely to be collected by liver, while hMSCs with moderate deformability were more likely in spleen but less in liver, although both these two groups of cells showed least preference to lung. It is suggested that the destination of cell microcirculation could be regulated by tuning cell mechanics finely, which is further discussed as follow.

Physiologically, arterial blood is pumped out from cardiopulmonary system, directly delivered to liver and spleen via artery blood vessels. Of note, venous blood output from spleen is next delivered to liver via portal vein. Portal vein blood from the other organs composes 75% of whole blood that a liver receives. It implies that liver can further collect the microcirculation cells escaping from spleen, besides these direct-supplied through heart-liver arterial blood vessels. When comparing with the results in present study, we can infer that HD-hMSCs could bypass spleen more easily than MD-hMSCs, due to their stronger deformability; hence, HD-hMSCs showed more human DNA in liver rather than in spleen. In addition, MD-hMSCs might be trapped in spleen due to their relatively weaker deformability before they are collected by liver. Hence, MD-hMSCs showed noticeable increased DNA in spleen compared with HD-hMSCs. Therefore, LD-cells, MD-cell, and HD-cells may together serve as a toolkit for customizing the microcirculation destination when passing lung-spleen-liver loop. That can potentially be applied in cell therapies for treating lung, spleen, liver correspondingly in a more precise manner.

In summary, this study has modified the deformability of hMSCs by culturing cells within the 3D dextran hydrogels containing crosslinking molecules distributed in optimized degrees of heterogeneity. The deformability of MSCs in petri dishes, microcosmically homogeneous (HOM) and heterogeneous (HET) dextran hydrogels here demonstrates a progressive increase according to the AFM test, which complies with the results on expression of F-actin derived from flow cytometry and fluorescent imaging. It was demonstrated that the modified hMSCs here, possessing enhanced deformability, can traversed the pulmonary vasculature and infiltrated the spleen and liver tissues in mice. This resulted in improved microcirculation and holds potential for facilitating stem cell therapy via intravenous administration.

## 4. Materials and Methods

### 4.1 Cell preparation

hMSCs were supplied by Shenzhen Zhongjia Bio-medical Technology Co., Ltd. (Guangdong Province, China). The culture medium comprised 89% (v/v) DMEM (Cat No: SH30023.01, HyClone, Logan UT, USA), 10% (v/v) fetal bovine serum (Cat No: 13011-8611, Every Green, Zhejiang Tianhang Biotechnology Co., Ltd, China), and 1% (v/v) penicillin-streptomycin (Cat No: SV30010, HyClone, Logan UT, USA). Cell incubator settings include 5% CO_2_ atmosphere and 37 °C temperature. Passage of used cells ranged from P3 to P9. To harvest hMSCs, the cells were digested with 0.09% trypsin (Cat No: SH30042.01, HyClone) in phosphate-buffered saline (PBS; Cat No: SH30256.01, HyClone) at 37 °C for 3.3 min. Next, the same volume of culture medium was added to neutralize the trypsin. Cells were then gently detached, centrifuged at 250×g for 6 min, and re-suspended in fresh medium for subsequent use.

### 4.2 Hydrogel fabrication

Maleimide-functionalized dextran (Mn = 42 kDa, Mw = 97 kDa, substitution degree = 6.32) (Cat No: FG91-1, Cellendes, Reutlingen, Germany) was used in this study. The maleimide-dextran was modified with cell-adhesive (RGD) peptides (Cat No: 09-P-001) via thiol-Michael addition, to obtain the cell-adhesive dextran. The reaction was conducted at room temperature for 10 min. The full sequence of RGD peptides is Acetyl-Cys-Doa-Doa-Gly-Arg-Gly-Asp-Ser-Pro-NH_2_, where Doa represents 8-amino-3,6-dioxaoctanoic acid. The reactive maleimide moieties of RGD-modified dextran were then blocked using monothioglycerol (Cat No: T10-3), also via thiol-Michael addition. The blocking reaction was conducted at room temperature for 10 min via the maleimide™thiol reaction. The designed final concentration for each component is indicated in Table 1.

Two equal-volume portions of a dextran solution were employed to create microscopic heterogeneity in the hydrogel composition. The final concentrations of the components in Part 1 and Part 2 are detailed in Table 1. The fabrication process of the three-dimensional(3D) dextran heterogeneous hydrogels is depicted in Figure 2A. Briefly, two aliquots of maleimide-dextran, each functionalized with varying amounts of available cross-linking sites, were thoroughly mixed and fully cross-linked. In the semi-synthesized hydrogel, this approach enhances the likelihood that two spatially adjacent dextran macromolecules will form covalent bonds with markedly different quantities of the cross-linker, a phenomenon referred to as the ‘cross-linker clustering (CLC) effect’.

The acidic buffer was added to control the final gelation pH to around 5.9. The pH was adjusted with HCl. reducing the reaction rate substantially, leaving enough time for the reaction components to fully mix. Next, pure culture medium (for hydrogel characterization) or the cell suspension was added and mixed to obtain the precursor solution. The dithiol cross-linker was then mixed with the precursor solution. The maleimide-dextran was cross-linked via thiol-Michael addition, and the hydrogel was formed after 5 min at room temperature. The maleimide-dextran, cross-linker, and buffer were all included in the same kit. The dithiol cross-linker consists of polyethylene glycol and the metalloprotease cleavable peptides (Pro-Leu-Gly-Leu-Trp-Ala) which can be cut by the metalloprotease secreted by cells inside hydrogels. The components of the 10-fold concentrated buffer included glucose (10gL^-1^), 2-(N-morpholino) ethanesulfonic acid (0.5 M), KCl (0.05 M), NaCl (1.1 M), NaH_2_PO_4_ (0.2 M), and phenol red (0.2 gL^-1^). The detailed information on the preparation for the working solution of each components of the HOM and HET dextran hydrogels are shown in the Section S1.1 of the Supplementary Materials.

### 4.3 Rheological measurement for hydrogels

The elastic modulus (G′) and the viscous modulus (G″) of 3D cross-linker-clustered dextran hydrogels were measured in a plate-to-plate rotational rheometer (Kinexus Pro, Malvern, U.K.). The samples were tested under 1% constant strain at 37 °C with frequency ranging from 0.01 Hz to 10 Hz. The gap distance was set at 0.2 mm. 90 µl hydrogel without cells was used for each test.

### 4.4 Cell deformability modification

10 µL hydrogel was fabricated in one well of 96-well plate for 3D culture of hMSCs, supplemented with 135 µL fresh medium. The medium was refreshed 1 hour later to wash out the residual components of the buffer. After that, the culture medium was refreshed every 2 days. Three hydrogel samples were prepared in the same reacting tube for each test group in one batch, and it was trisected to fabricate three samples of 10 µL hydrogel. The volume of cell suspension and hydrogel components used for fabricating 30 µL (3 × 10 µL) hydrogel was specified in Table S1. All the cell samples were observed by an inverted microscope (Olympus IX73, Olympus Corporation, Tokyo, Japan). Optical images were captured by a digital sCMOS camera (HAMAMATSU C11440-42U30, Hamamatsu Photonics K.K., Hamamatsu, Japan). That optical microscopy was also applied to image hMSCs in the subsequent characterizations.

### 4.5 Atomic force microscope (AFM) testing

The hMSCs were cultured in the context of 3D hydrogels or TCPS for 3 days. The hMSCs in hydrogels were then isolated by degrading dextran with a 1:20 (v/v) solution of dextranase (Cat No: D10-1, Cellendes, Reutlingen, Germany) diluted in culture medium, at 37 °C for 1 hour; and cells on TCPS were detached with trypsin. The isolated hMSCs were re-suspended with fresh culture medium, and transferred into a chamber slide (Cat No: 154534, Thermo Fisher Scientific, Waltham, MA, USA). hMSCs were cultured for 24 h and then fixed with 4% paraformaldehyde (PFA) in PBS for 25 min. Then, the chamber was removed, and the hMSCs on the slide were loaded directly for the AFM test. Young’s modulus was measured in the liquid phase by immersing hMSCs in PBS. A tip with a 10–20 kHz resonant frequency, 0.01–0.06 N m^-1^ spring constant, 220–230 µm length, and 15–25 µm width was used for the AFM (JPK NanoWizard III, Bruker Corp., Billerica, MA, USA). The software settings for the AFM test included a 0.5 nN setpoint, 10 µm z-length, and 2 µm s^-1^ loading and unloading speed. The vertical deflection was measured during the extended or retracted tip against the cell. Young’s modulus was calculated from the measured force-position curve during loading process using the JPK SPM data processing software, based on the Hertz model.

### 4.6 Immunofluorescence staining and microscopy

The sample was washed twice with PBS at room temperature for 5 min each time. Next, the sample was fixed with 4% paraformaldehyde (Cat No: 28908, Thermo Fischer Scientific, Waltham, MA, USA) in PBS for 25 min. The sample was then washed with PBS three times for 10 min each time. After fixation, the samples were permeabilized with 0.1% Triton X-100 (Cat No: X100-5ML, Sigma-Aldrich, St. Louis, MO, USA) in PBS for 7 min. The sample was then washed three times with PBS for 10 min each time.

Alexa Fluor 488-conjugated phalloidin (165 nM) (Cat No: A12379, Invitrogen, Carlsbad CA, USA) was dissolved in PBS to obtain the working solution for staining cellular F-actin. 1% (v/v) bovine serum albumin (Cat No: 37525, Thermo Fischer Scientific, Waltham, MA, USA) was added to the working solution to reduce the nonspecific background. The fixed and permeabilized samples were stained in the dark at room temperature for 1 h. Samples were then washed three times with PBS for at least 30 min each time.

DAPI (300 nM) (Cat No: A12379, Invitrogen, Carlsbad, CA, USA) was dissolved in PBS to obtain the working solution of nuclear staining. The fixed and permeabilized samples were stained in the dark at room temperature for 15 min, and then washed thrice with PBS for 15 min each time.

Fluorescent images were captured with an inverted microscope and a digital sCMOS camera. The regions of hMSCs in these images were recognized, and the fluorescent intensity of each stained protein in these regions was analyzed with the ImageJ software (National Institutes of Health, Bethesda, MD, USA). That result was used to measure the expression of the corresponding cellular protein.

### 4.7 Flow cytometry (FCM)

The cell suspension, obtained from the 3D hydrogels with microcosmic inner structure of homogeneous (HOM) and heterogeneity (HET) characteristics were quantitatively analyzed by FCM (BD FACSCanto II, BD Biosciences). The materials and methods of dissolving the 3D dextran hydrogels are shown in the Section S1.2 of the Supplementary Materials. The labeled F-actin proteins in cells were excited at the wavelength of 488 nm; and the corresponding emitted light was detected at 519 nm. More than 9 × 10^3^ cells were analyzed in each sample run. The statistic of the FCM data were processed in FlowJo software (FlowJo v10, Becton, Dickinson & Company, trial version).

### 4.8 Animal experiment

The hMSCs cultured in hydrogels for 3 days were isolated by degrading dextran with a 1:20 (v/v) diluted solution of dextranase (Cat No: D10-1, Cellendes, Reutlingen, Germany) in culture medium, at 37 °C for 1 hour; and cells cultured on TCPS for 3 days were detached with trypsin solution. The isolated hMSCs were resuspended with fresh culture medium for the following animal experiment. The hMSCs were collected by centrifuging and resuspended again in sterile saline. 200 µL normal sterile saline carrying 1 × 10^5^ hMSCs was injected into each nude mouse (Balb/c, Changzhou Cavens Experimental Animal Co., Ltd, China) through the tail vain. Organs including the liver, spleen and lung of the mice were taken after 20 min. The blood remaining in the atria and ventricles of the heart was removed. Approximately 50 mg of tissue was sectioned from each organ for PCR test. The hMSCs modified in 3D hydrogels or on the TCPS were tested. 3 mice were sacrificed in each test group, and 4 replications of RT-qPCR experiment were conducted on each organ tissue taken from each mouse.

### 4.9 RT-qPCR

Human Alu sequences were used to verify the existence of hMSCs in the mice. Housekeeping gene of mice, GAPDH, was selected as the reference. Total genomic deoxyribonucleic acid (DNA) was isolated by TRI reagent (Cat No: T9424, Sigma-Aldrich, USA) according to the standard protocol provided by the manufacturer. The reaction solution for qPCR was composed of taqman PCR mix (Cat No: SR2110, Solarbio, China), 0.2 µM each of forward and reverse primers, 0.2 µM Alu probe and approximately 10 µg DNA for the detection of Alu sequences or 100 ng DNA for GAPDH. The polymerase activation and DNA denaturation were performed at 50 °C for 2 min and 95 °C for 10 min. Amplification progressed 40 cycles, each of which including denaturation at 95 °C for 15 s and annealing with extension at 60 °C for 1 min. The melting curve was obtained by protocols demonstrated by the manufacturer (LightCycler 480 II, Roche, Switzerland). The relative level of the human DNA was quantified by calculating the formula 2^-ΔCt^. ΔCt represents the threshold cycle of the human Alu sequences subtracted by the mice GAPDH. The corresponding sequences were listed in Table 2.

**Table 2.**
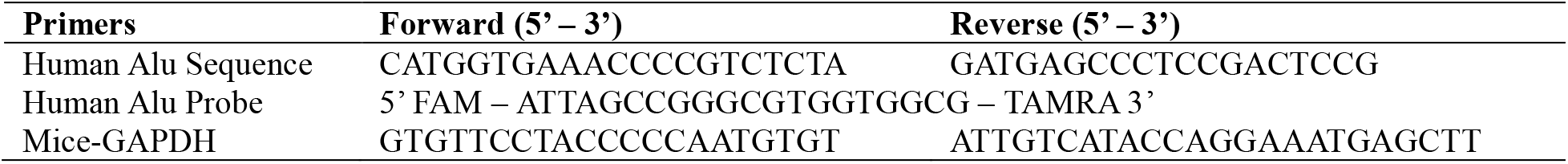
Sequences of primers and probe used for RT-qPCR.

### 4.10 Tissue sample preparation for transmission electron microscopy

The tissue samples were cut and modified into <1mm’ small pieces. After centrifugation and precipitation, the cell samples were fixed with pre-cooled glutaraldehyde and placed at 4°C for later use. The sample was washed 4 times with phosphoric acid buffer for 1 hour each time. Then it was fixed with 1% osmic acid at 4°C for 2 hours, and then washed three times with ddH_2_O for 10 minutes each time; Then it was stained in 2% uranium acetate aqueous solution for 2 hours, followed by the dehydration process using alcohol with gradient concentration, and then infiltrated with the mixed 100% acetone and EPON812 resin with a ratio of 1:1 for 2 hours (room temperature), and then infiltrated with pure EPON812 resin (embedding agent) in the embedding mold overnight at 30 °C. After that, the polymerization was carried out in a 37°C incubator for 12 hours. Subsequently, the ultrathin sections were made by Leica EM UC7 micro-slicing machine with a thickness of about 70nm. After staining with lead nitrate for 5 minutes and washing for 3 times, the ultrathin sections were observed by electron microscopy (electron microscope model FEI Tecnai G2 120kV).

### 4.11 Statistical analysis

Data are presented as the mean ± standard deviation (SD). Statistical significance was assessed using the one-way analysis of variance (ANOVA) with Tukey’s post-hoc analysis. All experiments were repeated at least three times.

## Acknowledgements

This work was funded by the National Natural Science Foundation of China (51875170), Fundamental Research Funds for the Central Universities of China (B200202225), Changzhou Sci&Tech Program (CE20195037). The authors thank Changzhou University for providing the rotational rheometer, and thank Dr. Xiuli Cong from Zhejiang hospital for gifting the cells and discussion for the cell culture technology and research result. The authors also thank the Nuo Ye from Nanjing maternity and child health care hospital for assisting the animal experiment.

## Author contributions

X. Z., Z. W., F. H. and F. T. performed the synthesis and characterization of hydrogel material, 3D cell culture experiment, FCM, AFM and TEM tests. X. Z., Z. W. and F. H. conducted the cellular staining, microscopy and PCR work. Y. S. and S. Y. conducted the mouse experiments, X.Z., Z. W., F. T. and Y. S. performed the data analysis, X.Z, Z. W. and F. T. conceived the project and designed the experiments. X.Z and F. T. supervised the project. X. Z. and Z.W. wrote the paper, and all authors contributed to the final draft.

## References

1. M. L. Skelton et al., Modular Multiwell Viscoelastic Hydrogel Platform for Two- and Three-Dimensional Cell Culture Applications. ACS Biomater Sci Eng 10, 3280–3292 (2024).

2. S. Wu et al., Regulating the migration of smooth muscle cells by a vertically distributed poly (2-hydroxyethyl methacrylate) gradient on polymer brushes covalently immobilized with RGD peptides. Acta biomaterialia 75, 75–92 (2018).

3. X. Yin, X. Zhu, Z. Wang, Cell Migration Regulated by Spatially Controlled Sti ness inside Composition-Tunable ThreeDimensional Dextran Hydrogels. Advanced Materials Interfaces 10.1002/admi.202100494 (2021).

4. J. Zhang et al., Dynamic Mechanics-Modulated Hydrogels to Regulate the Differentiation of Stem-Cell Spheroids in Soft Microniches and Modeling of the Nonlinear Behavior. Small 15, 1901920 (2019).

5. S. H. Hsu, P. S. Hsieh, Self-assembled adult adipose-derived stem cell spheroids combined with biomaterials promote wound healing in a rat skin repair model. Wound Repair and Regeneration 23, 57–64 (2015).

6. C. Gwam, N. Mohammed, X. Ma, Stem cell secretome, regeneration, and clinical translation: a narrative review. Annals of Translational Medicine 9 (2021).

7. J. A. Brassard, M. Nikolaev, T. Huebscher, M. Hofer, M. P. Lutolf, Recapitulating macro-scale tissue self-organization through organoid bioprinting. Nature Materials 20, 22–29 (2021).

8. A. Lewis, R. Keshara, Y. H. Kim, A. Grapin-Botton, Self-organization of organoids from endoderm-derived cells. Journal of Molecular Medicine 99, 449–462 (2021).

9. X. Zhu, Z. Wang, F. Teng, A review of regulated self-organizing approaches for tissue regeneration. Progress in Biophysics and Molecular Biology 10.1016/j.pbiomolbio.2021.07.006 (2021).

10. Z. Fan et al., Acoustic Actuation of Integrin‐Bound Microbubbles for Mechanical Phenotyping during Differentiation and Morphogenesis of Human Embryonic Stem Cells. Small 14, 1803137 (2018).

11. C. Lin et al., Matrix promote mesenchymal stromal cell migration with improved deformation via nuclear stiffness decrease. Biomaterials 217, 119300 (2019).

12. G. Nardone et al., YAP regulates cell mechanics by controlling focal adhesion assembly. Nature Communications 8 (2017).

13. A. Tajik et al., Transcription upregulation via force-induced direct stretching of chromatin. Nature materials 15, 1287 (2016).

14. K. Ghosh et al., Cell adaptation to a physiologically relevant ECM mimic with different viscoelastic properties. Biomaterials 28, 671–679 (2007).

15. M. E. Fernandez-Sanchez, T. Brunet, J. C. Röper, E. Farge, Mechanotransduction’s impact on animal development, evolution, and tumorigenesis. Annu Rev Cell Dev Biol 31, 373–397 (2015).

16. W. Sang, A. Ural, Quantifying how altered lacunar morphology and perilacunar tissue properties influence local mechanical environment of osteocyte lacunae using finite element modeling. J Mech Behav Biomed Mater 135, 105433 (2022).

17. S. K. Venkatesh, M. Yin, R. L. Ehman, Magnetic resonance elastography of liver: Technique, analysis, and clinical applications. Journal of Magnetic Resonance Imaging 37, 544–555 (2013).

18. A. Massey et al., Mechanical properties of human tumour tissues and their implications for cancer development. Nat Rev Phys 6, 269–282 (2024).

19. J. Cooper, F. G. Giancotti, Integrin Signaling in Cancer: Mechanotransduction, Stemness, Epithelial Plasticity, and Therapeutic Resistance. Cancer Cell 35, 347–367 (2019).

20. B. A. Radman, A. M. M. Alhameed, G. Shu, G. Yin, M. Wang, Cellular elasticity in cancer: a review of altered biomechanical features. J Mater Chem B 12, 5299–5324 (2024).

21. J. Wei, M. Li, Interplay of Fluid Mechanics and Matrix Stiffness in Tuning the Mechanical Behaviors of Single Cells Probed by Atomic Force Microscopy. Langmuir 10.1021/acs.langmuir.2c03137 (2023).

22. N. O. Chahine et al., Effect of age and cytoskeletal elements on the indentation-dependent mechanical properties of chondrocytes. PloS one 8, e61651 (2013).

23. L. R. Smith, S. Cho, D. E. Discher, Stem Cell Differentiation is Regulated by Extracellular Matrix Mechanics. Physiology (Bethesda) 33, 16–25 (2018).

24. L. Xiao, Y. Sun, L. Liao, X. Su, Response of mesenchymal stem cells to surface topography of scaffolds and the underlying mechanisms. J Mater Chem B 11, 2550–2567 (2023).

25. D. M. Graham, K. Burridge, Mechanotransduction and nuclear function. Current opinion in cell biology 40, 98–105 (2016).

26. S. Tietze et al., Spheroid culture of mesenchymal stromal cells results in morphorheological properties appropriate for improved microcirculation. Advanced Science 6, 1802104 (2019).

27. M. Sanchez-Diaz et al., Biodistribution of Mesenchymal Stromal Cells after Administration in Animal Models and Humans: A Systematic Review. Journal of Clinical Medicine 10, 2925 (2021).

28. S.-j. Kim et al., Spatially Arranged Encapsulation of Stem Cell Spheroids Within Hydrogels for Regulation of Spheroid Fusion and Cell Migration. SSRN Electronic Journal (2021).

29. J. Park, G. Choe, S. Oh, J. Y. Lee, In Situ Formation of Proangiogenic Mesenchymal Stem Cell Spheroids in Hyaluronic Acid/Alginate Core/Shell Microcapsules. EngRN: Tissue Engineering (Topic) (2020).

30. Z. Wang, X. Zhu, X. Cong, Spatial micro-variation of 3D hydrogel stiffness regulates the biomechanical properties of hMSCs. Biofabrication 13, 035051 (2021).

31. K. Funakoshi et al., Highly sensitive and specific Alu-based quantification of human cells among rodent cells. Scientific reports 7, 1–12 (2017).

