## Supplemental materials for "Modulating Cellular Deformability via 3D Dextran Hydrogel Cultivation to Regulate the Microcirculation of Mesenchymal Stem Cells in Murine Spleen and Liver"

### S1 Supplementary Materials and Methods

#### S1.1 Hydrogel components

Ten-fold concentrated buffer (10× CB) is used to adjust the pH value of the fabrication environment. The 3D biodegradable cross-linker-clustered MD hydrogel was fabricated at around pH 5.9 because a slightly acidic environment could reduce the reaction rate substantially, leaving enough time for complete mixing of the reaction components. Deionized water is used to keep the final volume of hydrogels constant in each case.

The components of the buffer include glucose (10 g/L), 2-(N-morpholino) ethanesulfonic acid (0.5 M), KCl (0.05 M), NaCl (1.1 M), $NaH_2PO_4$ (0.2 M) and phenol red (0.2 g/L); and the pH is adjusted with HCl. For the reaction components of the hydrogel, thiol groups are determined with the Ellman assay, which measures appearance of a yellow color up on the thiol-induced cleavage of the disulfide bond of 5,5'-dithiobis-(2-nitrobenzoic acid). Maleimide groups are measured by using their light absorption at 310 nm under UV spectroscopy. There is 30 mM maleimide in dextran working solution, 20 mM thiol in CD-Link working solution, 0.3 mM thiol in RGD peptide working solution, and 3 mM thiol in thioglycerol working solution (Table S1). The volume of each working solution is listed for the HOM and HET hydrogel, respectively, as shown in Table S1.

Table S1. Volume of cell suspension and hydrogel components.

| **Cross-Linker-Clustering Degree** | | | **HOM** | **HET** |
|---|---|---|---|---|
| Precursor Solution | Part 1 | Maleimide Groups of Dextran (30 mM maleimide groups) [µL] | 1.5 | 1.5 |
| | | 1-Thioglycerol (3 mM thiol groups) [µL] | 4.8 | 6.7 |
| | Part 2 | Maleimide Groups of Dextran [µL] | 1.5 | 1.5 |
| | | 1-Thioglycerol (3 mM thiol groups) [µL] | 4.8 | 2.9 |
| | Arg-Gly-Asp Peptide (Single-Thiol Modified RGD, 0.3 mM thiol groups) [µL] | | 4 | 4 |
| | Cells Suspension (1.2 × $10^4$ cells/µL) [µL] | | 5 | 5 |
| | 10× Buffer | | 1× | 1× |
| | DI water [µL] | | 2.9 | 2.9 |
| Thiol Groups within the Cross-Linker (Dithiol Modified, 20 mM thiol groups) [µL] | | | 2 | 2 |
| Total Volume [µL] | | | 30 | 30 |

#### S1.2 Hydrogel dissolving for flow cytometry

The culture medium was sucked out from the microwell. The sample was cut into pieces with the suction head of pipette, covered with a 1:15 (v/v) diluted solution of dextranase (Cat no.: D10-1, Cellendes, Reutlingen, Germany) in culture medium, and incubated at 37 °C for 30 min in a standard $CO_2$ incubator. After the gel being dissolved, the cells can be observed to suspend in the medium and can be shaken with the fluidic medium. The resulting cell suspension in 1.5 mL tube was centrifuged at 1200 rpm for 4 min, and the cells were resuspended in PBS. The dextranase used here was not reacting with cells, not harmful for cells and able to keep the cells alive when dissolving the dextran hydrogels.

#### S2. Supplementary Figures

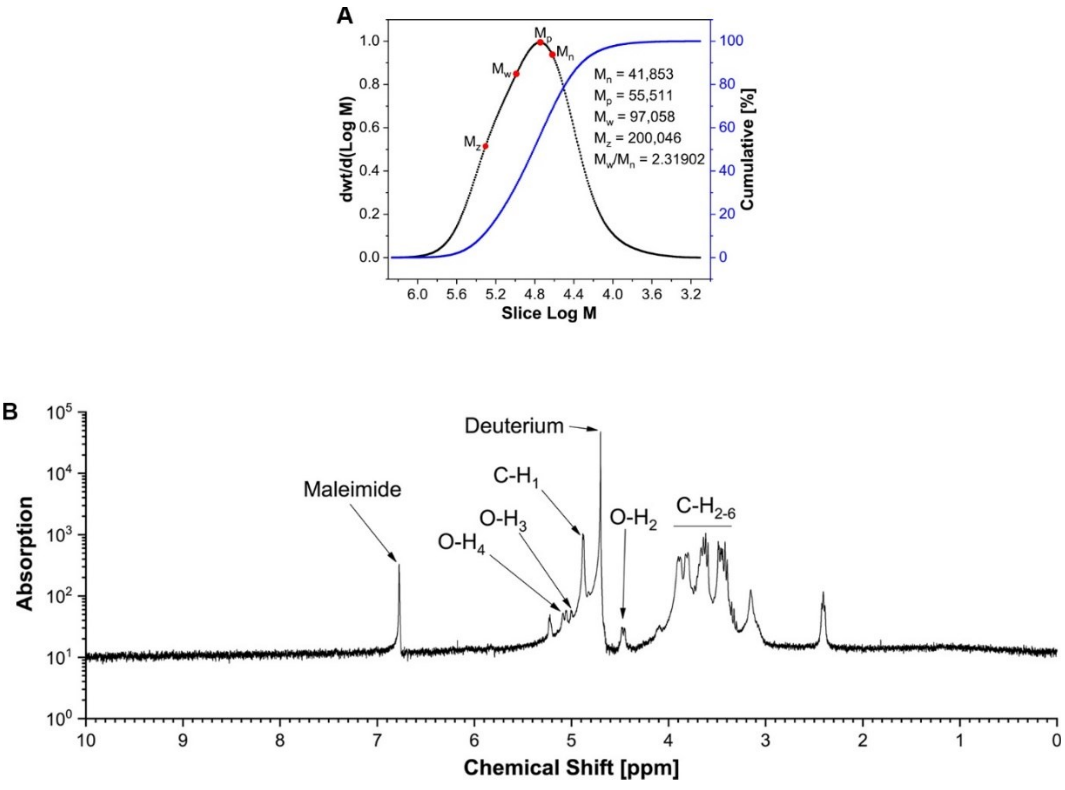


**Figure S1.** Characterization of maleimide-dextran polymer. (A) Molecular weight distribution of maleimide-functionalized dextran polymer. (B) $^1$H NMR spectra of the maleimide-dextran in deuterium oxide under 400 MHz.
